# Likelihood Ratios Given Activity-Level Propositions for DNA Transfer Evidence: Practical Implementation and Simulation Studies Using the HaloGen Engine (Part II)

**DOI:** 10.64898/2026.02.06.703509

**Authors:** Peter Gill, Øyvind Bleka

**Affiliations:** Department of Forensic Sciences, Oslo University Hospital, Oslo, Norway; Department of Clinical Medicine, University of Oslo, Oslo, Norway

## Abstract

The quantitative interpretation of low-template DNA findings given activity-level propositions requires models that can accommodate inter-laboratory variability, uncertainty in transfer and recovery, and case-specific assumptions. This paper presents the practical implementation of `HaloGen`, an open-source hierarchical Bayesian framework for calculating activity-level likelihood ratios (LRs) from DNA quantity data.

We compare three modelling approaches derived from the framework: a *Group* model, which combines data across laboratories, a hierarchically informed *Lab–Bayes* model, and a standalone, laboratory specific *Lab–Vague* model. Through simulation studies, we show that evidential strength is sensitive not only to DNA quantity but also to case context, particularly the assumed number of relevant actors (*N*_*S*_), the treatment of specified unknown contributors, and the choice of laboratory calibration. Inter-laboratory differences in DNA recovery and non-detection can lead to materially different LRs when these data are used within the HaloGen framework, so pooled or external data should not be used uncritically.

To address practical implementation, we propose a minimum-effort calibration pathway for laboratories wishing to use `HaloGen` for quantitative activity-level LR reporting. The results indicate that a limited number of local direct/secondary transfer experiments can improve relevance compared with exclusive reliance on a pooled population model, although the adequacy of any dataset remains case- and proposition-dependent. The findings clarify how contextual assumptions enter mathematically into activity-level inference and underscore the importance of transparent specification of propositions, data relevance, model assumptions, and remaining expert judgement.

## 1. Introduction

The practical evaluation of low-template DNA evidence recovered from touched items is challenging due to substantial variability in DNA transfer, persistence, and recovery, arising from differences in substrates, activities, and laboratory procedures [1]. While the likelihood ratio (LR) provides a logically sound framework for evaluating biological results under activity-level propositions [2], its application depends critically on the availability of probabilistic models that are both robust to uncertainty and adaptable to case-specific context.

The ReAct project demonstrated, on a large scale, the extent of inter-laboratory variability in experimental DNA transfer data. Building on this, Taylor et al. [3] proposed a Bayesian network approach in which such variability was incorporated as a post-hoc adjustment factor, providing a pragmatic strategy when laboratories must rely on external data. While conceptually important, this approach does not directly address how laboratory-specific performance, uncertainty, and case assumptions should be integrated into a unified probabilistic framework.

The present study advances this line of work by adopting hierarchical Bayesian models, which allow laboratory-specific parameter estimation while borrowing strength from cross-laboratory data [4]. This structure provides a principled mechanism for stabilising inference in data-limited settings while preserving genuine differences between laboratories.

This paper is the second in a two-part series. In Part 1 [5], we developed the theoretical foundations of the HaloGen (Hierarchical Bayesian Activity Level Orchestrator: next Generation) framework, including the mathematical formulation of activity-level likelihood ratios for scenarios involving multiple contributors, multiple relevant actors, and multiple stains.

Here, in Part 2, we focus on the practical implementation and application of HaloGen using an open-source R framework. We examine how evidential conclusions depend on modelling choices by comparing likelihood ratios obtained under three approaches that arise naturally from the hierarchical structure:

1. A *Group model* that pools data across laboratories;
2. A hierarchically informed *lab-specific (Lab–Bayes) model*, which uses the *Group* model to form informative priors and represents the recommended approach; and
3. A *standalone lab-specific (Lab–Vague) model*, which relies primarily on data from a single laboratory under weakly informative priors.

Through a series of simulation studies, we investigate how likelihood ratios respond to changes in DNA quantity, laboratory performance, numbers of contributors, numbers of actors, and the amount of available calibration data. In doing so, we show that contextual assumptions, particularly those relating to actor number and background or specified unknown contributors, inform the LR calculation. This also provides a formal basis for understanding how confirmation bias may arise from unexamined modelling choices: throughout the results we show that fixing contextual assumptions such as the number of actors (*N*_*S*_), the treatment of unknown contributors, or the use of pooled versus lab-specific parameters can systematically inflate or deflate likelihood ratios even when the observed DNA quantities are unchanged. This highlights the importance of transparent specification of propositions and parameters, and of examining plausible alternative assumptions rather than conditioning implicitly on a single favoured scenario.

The purpose of this paper is not to suggest that all activity-level opinions must be generated by HaloGen or by any single statistical model. Activity-level evaluation may also involve probability assignments informed by experimental data, published literature, case-specific information, or expert judgement, provided that the basis, assumptions, and limitations of those assignments are made explicit. The results presented here concern quantitative, model-based LR reporting within the `HaloGen` framework and the ReAct-type direct/secondary transfer data analysed in this paper.

The remainder of the paper describes the simulation design and modelling architecture, followed by a combined Results and Discussion section presenting the experimental findings and their implications for forensic practice. We conclude by discussing validation requirements, practical implementation, and the broader requirement for robust and balanced activity-level interpretation.

## 2. Materials and Methods

### 2.1. Notation

All likelihood ratios (LRs) reported in this paper are Bayes factors comparing the probability of the observed data *E* under competing activity-level propositions. Elemental hypotheses are assigned equal prior probabilities, following the approach described in Part 1 [5] (Section 2.4), modified from [6]. For notational simplicity, we use the LR symbol throughout; for example, log_10_LR_case,S1_ denotes the case-level LR for suspect 1 on a logarithmic scale (base 10). A complete list of symbols and notation is provided in Supplement S1.

### 2.2. Experimental Data

The experimental data analysed in this study are described in detail in [5] (Section 3.8). A total of 20 sets of laboratory results were used to construct and evaluate the models. Direct and secondary transfer experiments were conducted one hour after hand shaking followed by handling a screwdriver handle.

### 2.3. Model Framework

The HaloGen framework is based on the zero-augmented, left-censored transfer model described in Part I [5]. For a transfer pathway with parameters *θ* = (*µ, σ, k*), the recovered DNA quantity *Q* is modelled as a mixture of a zero-transfer/no-transfer component and a positive lognormal transfer component. The parameter *k* denotes the probability that no measurable DNA is transferred, while *µ* and *σ* parameterise the lognormal distribution of positive quantities.

A recorded non-detect can therefore arise in two ways: either no measurable DNA is transferred, with probability *k*, or a positive transfer occurs but falls below the detection limit DL. Thus,

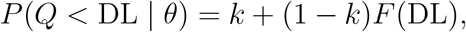

where *F* (DL) is the lognormal distribution function evaluated at the detection limit. Equivalently, for the lognormal component,

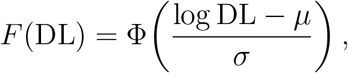

where Φ(·) is the standard normal distribution function.

For an observed detected experimental quantity *q*_obs_ ≥ DL, the contribution used when fitting the transfer model is the continuous density

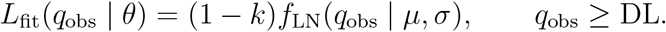

Thus, in the experimental fitting likelihood, non-detects contribute through probability mass, whereas observed detected quantities contribute through a density.

At case level, a further conditioning step is used. Once a contributor quantity *q* ≥ DL has been observed, detection has already occurred, and the case-level likelihood contribution is evaluated using the density of the observed quantity conditional on detection:

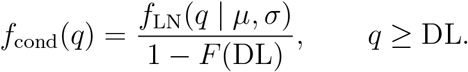

This conditioning rule is applied to both the direct-transfer and secondary-transfer pathways. We write

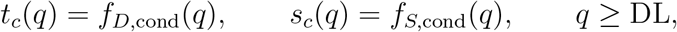

for the corresponding conditional-on-detection likelihood components for contributor *c*, under the direct-transfer and secondary-transfer pathways, respectively.

Non-detects are handled separately through probability-mass terms, rather than as zero-valued density observations. In particular, where an elemental hypothesis requires a relevant actor who is not represented among the detected contributors, HaloGen uses the actor non-detection probability *F*_0_. This aligns the implementation used here with the theoretical definitions in Part I.

The fail-rate parameter *F*_0_ denotes the probability that a relevant actor leaves no detectable DNA:

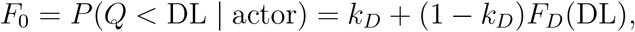

where the subscript *D* denotes the direct-transfer model. In the current implementation, *F*_0_ is estimated from direct-transfer non-detection information using a Jeffreys-smoothed empirical Beta policy with clamping, as described in Part I [5].

Each laboratory *L* has its own parameter set (*µ*_*L*_, *σ*_*L*_, *k*_*L*_), modelled hierarchically with group-level hyperparameters. In the cross-laboratory *Group* model these parameters are partially pooled across laboratories. In the *Lab–Bayes* model, the *Group* model informs laboratory-specific priors, while the *Lab–Vague* model estimates laboratory parameters using weakly informative priors and only the local laboratory data. Formal definitions of all statistical parameters, variables, likelihood components, the *F*_0_ policy, and the exhaustive hypothesis construction are provided in Part I [5].

### 2.4. Bayesian Inference

Posterior inference was performed using Markov chain Monte Carlo (MCMC) sampling implemented in `Stan` [7], interfaced via `R`. Four independent MCMC chains were run for each model. Convergence was assessed using Stan’s rank-normalised split-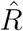 diagnostic and associated bulk and tail effective sample sizes, as implemented in Stan.

Likelihood ratios were computed on a per-draw basis using paired posterior samples, that is, the same posterior draw index was used consistently in both the numerator and denominator. The resulting distribution of log_10_(LR) values was summarised by its median, with 10–90% intervals reported to characterise uncertainty.

### 2.5. Simulation Design

All simulations use the exhaustive-proposition framework described in [5], conditioning on a specified number of relevant actors (*N*_*S*_). For example, when *N*_*S*_ = 1, exactly one actor is assumed to have performed the activity specified by the proposition. That actor may correspond to the specified suspect (*S*_1_), to a specified unknown contributor (e.g. *U*_1_), or to an unobserved actor who leaves no detectable DNA, represented through the fail-rate parameter *F*_0_. DNA may therefore be recovered from one or more specified contributors, or from none of the specified contributors, with all admissible possibilities incorporated explicitly into the hypothesis space.

For each posterior draw of the model parameters obtained via MCMC, HaloGen evaluates the full case-level likelihood under the competing activity-level propositions and computes a corresponding likelihood ratio. The resulting collection of per-draw LRs provides a Monte Carlo representation of the posterior-marginalised LR.

Simulation Experiments A–C, described in sections 3.7 and 3.8, were designed to examine model behaviour under progressively more realistic laboratory constraints, including limited laboratory-specific data and varying combinations of direct and secondary transfer experiments.

### 2.6. Priors and Detection-Limit Validation

All prior specifications follow those described in Supplement S4.1. Weakly informative priors were used for lab-level transfer parameters. Defence-conservative behaviour at the case level was ensured through an empirically calibrated fail-rate term *F*_0_, based on a Jeffreys-smoothed Beta(0.5, 0.5) prior whose posterior draws are bounded within a data-dependent interval to prevent unrealistically small or large values. This policy prevents anti-conservative likelihood ratios when non-detect rates are sparse.

To assess robustness to uncertainty in the analytical detection limit, a detection-limit sensitivity analysis was performed using the “DL robustness” test implemented in `Validation.R` (Supplement S4.1). Two plausible detection-limit thresholds (low and high) were analysed, and robustness was assessed by the absolute difference in log_10_(LR) between the two settings.

### 2.7. Validation and Diagnostics

All simulations were verified using the `gemini_lockdown.R` diagnostics suite. These checks verify consistency with the HaloGen engine, and confirm expected theoretical behaviour; for example, that the likelihood ratio does not decrease when the recovered DNA quantity from the person of interest increases while all other aspects of the case are held fixed. Unless otherwise stated, all analyses used the exhaustive *N*_*S*_ = 1 configuration and the empirical *F*_0_ policy described in [5].

### 2.8. Software and Resources

All programs, user manuals, and data used to generate the results in this paper are available at https://sites.google.com/view/altrap/halogen and https://github.com/peterdgill/HaloGen_Bayes. HaloGen_v2.0.2 was used.

## 3. Results and Discussion

This section presents simulation results illustrating how the HaloGen framework behaves under realistic activity-level scenarios. The emphasis is on identifying trends and practical implications for casework rather than exhaustively narrating numerical values, which are reported in the accompanying tables and figures.

### 3.1. Interpretation via Probability Density Functions

Figure 1 shows posterior predictive probability density functions for DNA quantities under direct and secondary transfer, comparing three modelling approaches: Group, Lab–Bayes, and Lab–Vague. Results are shown for a high-DNA-recovery laboratory (lab_1_ESS) and a low-DNA-recovery laboratory (lab_10). Across both laboratories, direct-transfer distributions are shifted toward higher quantities than secondary-transfer distributions. Differences between models reflect how information is pooled. The Group model represents an average laboratory and smooths over laboratory-specific performance. The Lab–Vague model reflects only local data and can exhibit wide or unstable tails when data are sparse. The Lab–Bayes model occupies an intermediate position, retaining laboratory specificity while moderating extreme behaviour through hierarchical pooling.

**Figure 1:**
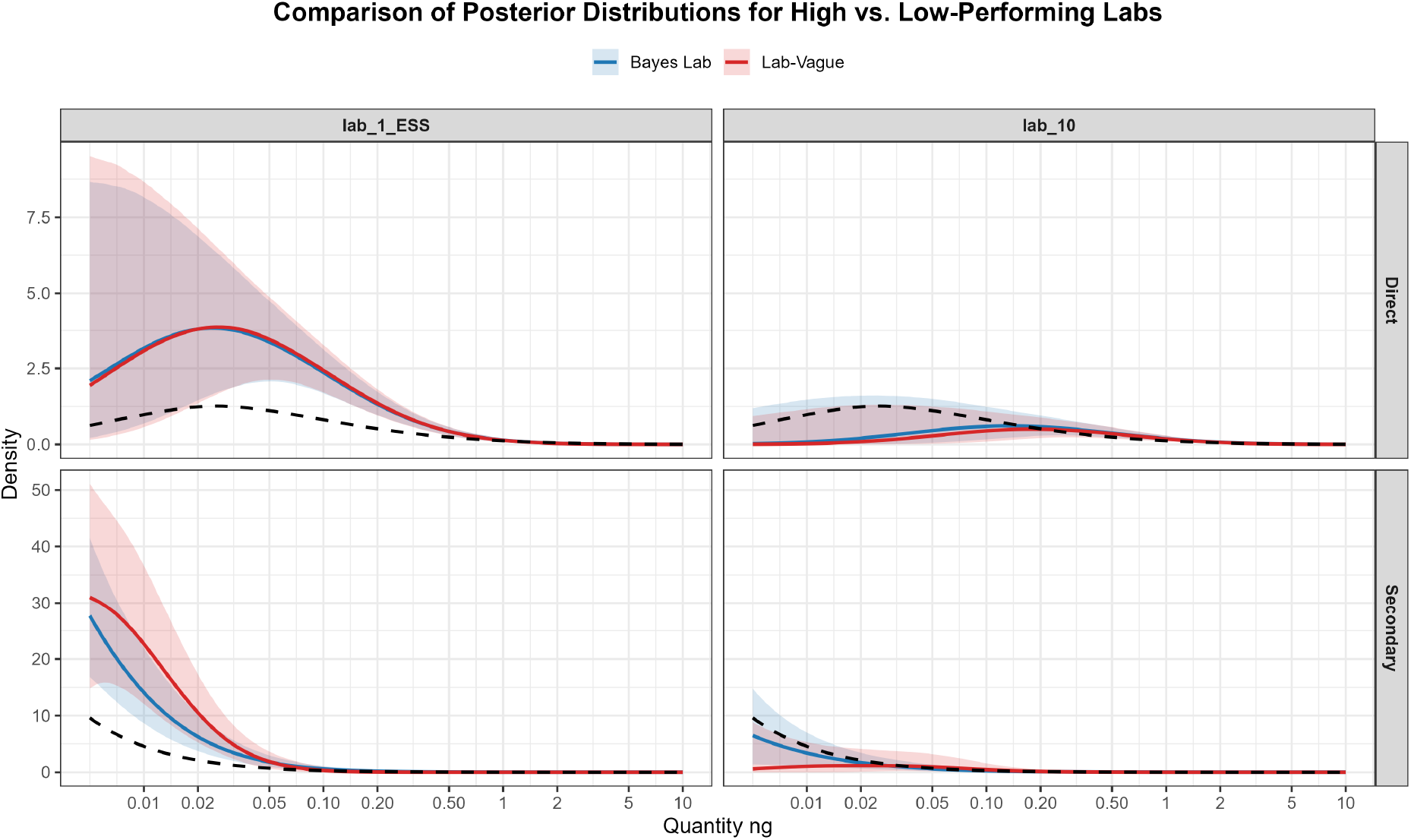
Posterior predictive probability density functions (PDFs) for Direct Transfer (top panels) and Secondary Transfer (bottom panels), plotted on a logarithmic scale. Red: Standalone *Lab–Vague* model; Blue: *Lab–Bayes* model; Black (dashed): *Group* model based on mean hyperparameters. The examples shown correspond to lab_1_ESS (high-DNA-recovery laboratory, left) and lab_10 (lower-DNA-recovery laboratory, right). Shaded regions represent 95% credible intervals derived from posterior draws. Differences between the curves reflect how model hierarchy and prior structure influence the inferred DNA quantity distributions.

Although these predictive distributions show substantial uncertainty in absolute quantities, the corresponding likelihood ratios are markedly more stable (Fig. 2). This occurs because likelihood ratios are formed from paired posterior draws: correlated uncertainty in direct and secondary transfer parameters largely cancels, yielding tighter distributions of log_10_(LR). This stabilisation is most pronounced under the Lab–Bayes model.

**Figure 2:**
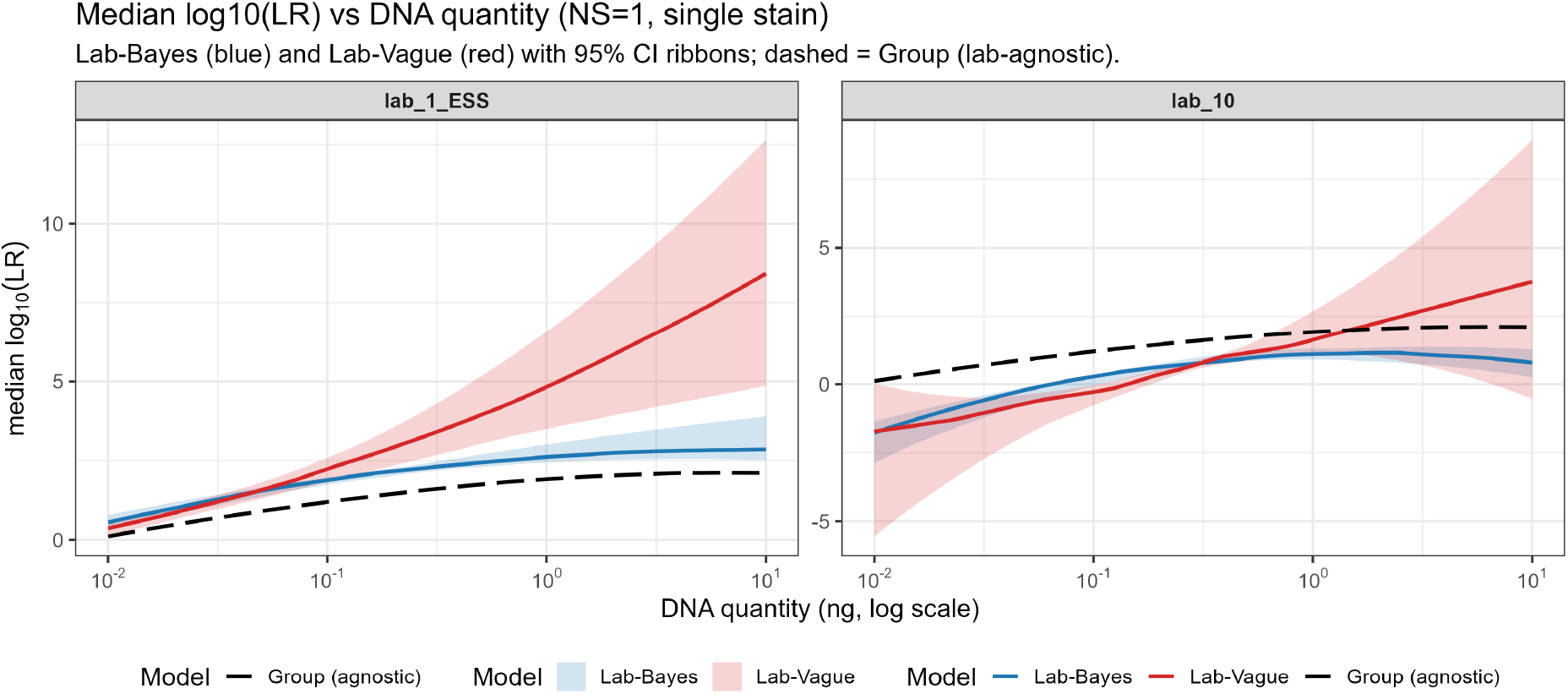
Median log_10_(LR) as a function of DNA quantity for high- and low-DNA-recovery laboratories. Blue: *Lab–Bayes* model; Red: *Lab–Vague* model; Black (dashed): *Group* model. Shaded ribbons represent 95% credible intervals across posterior draws. The *Lab–Vague* model shows greater sensitivity (wider intervals and occasional inflation) at higher DNA quantities, while the *Lab–Bayes* model maintains stable, conservative estimates.

### 3.2. Effect of DNA Quantity on Likelihood Ratios

Tables 1 and 2 summarise how the case-level LR for a single person of interest varies with recovered DNA quantity when a single relevant actor is assumed. An illustrative sensitivity analysis showing how the case-level LR depends jointly on DNA quantity, mixture composition, the number of contributors, and the assumed number of relevant actors (e.g. offenders) is provided in Supplement S5.

**Table 1:** High DNA recovery laboratory (lab_1_ESS): Effect of DNA quantity on log_10_ LR_case,*S*1_ for single-stain scenarios (*N*_*K*_ = 1, *N*_*U*_ = 0, *N*_*S*_ = 1). Medians shown for Group, Lab–Bayes, and Lab– Vague models.

| Case ID | $q_{S_1}$ (ng) | $\log_{10}(\text{LR})_{S_1}$ (Group) | $\log_{10}(\text{LR})_{S_1}$ (Lab-Bayes) | $\log_{10}(\text{LR})_{S_1}$ (Lab-Vague) |
| --- | --- | --- | --- | --- |
| Very Low Q | 0.01 | 0.09 | 0.61 | 0.42 |
| Low Q | 0.05 | 0.89 | 1.58 | 1.59 |
| Medium Q | 1.00 | 1.92 | 2.61 | 4.88 |
| High Q | 1.50 | 2.00 | 2.68 | 5.46 |
| Very High Q | 5.00 | 2.12 | 2.80 | 7.36 |

**Table 2:** Low DNA recovery laboratory (lab_10): Effect of DNA quantity on log_10_ LR_*S*1_ for single-stain scenarios (*N*_*K*_ = 1, *N*_*U*_ = 0, *N*_*S*_ = 1). Medians shown for Group, Lab–Bayes, and Lab–Vague models.

| Case ID | $q_{S1}$ (ng) | $\log_{10}(\text{LR})_{S1}$ (Group) | $\log_{10}(\text{LR})_{S1}$ (Lab-Bayes) | $\log_{10}(\text{LR})_{S1}$ (Lab-Vague) |
| --- | --- | --- | --- | --- |
| Very Low Q | 0.01 | 0.10 | -1.77 | -1.71 |
| Low Q | 0.05 | 0.89 | -0.27 | -0.75 |
| Medium Q | 1.00 | 1.92 | 1.11 | 1.61 |
| High Q | 1.50 | 2.01 | 1.15 | 1.97 |
| Very High Q | 5.00 | 2.11 | 1.11 | 3.11 |

For the high-DNA-recovery laboratory (lab_1_ESS), increasing DNA quantity produces a monotonic increase in evidential support for direct transfer. The most pronounced change occurs at low quantities, where the LR transitions from near-neutral to strong support. For the low-DNA-recovery laboratory (lab_10), low quantities tend to favour secondary transfer, and even higher quantities yield only modest support for direct transfer.

These results demonstrate that evidential strength is not determined by DNA quantity alone, but by quantity interpreted through laboratory-specific transfer characteristics. Group-based models can understate evidence for high-recovery laboratories and overstate it for low-recovery laboratories, whereas the Lab–Bayes model provides more balanced behaviour across conditions.

### 3.3. Extension to Multiple Stains

#### 3.3.1. Replicate Stains from a Single Contributor

Table 3 shows results for scenarios involving three replicate stains attributed to a single contributor. When all stains yield very low quantities, evidential support remains weak or favours secondary transfer. When all stains yield moderate or high quantities, evidential support increases substantially relative to a single-stain scenario, reflecting accumulation of consistent observations.

**Table 3:**
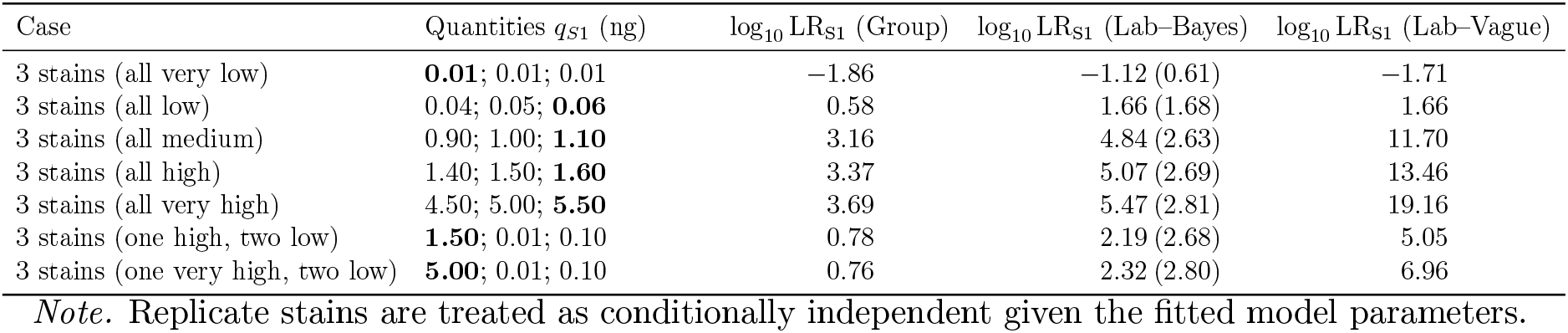
Lab_1_ESS: Effect of replicate stains for S1 on log_10_ LR_S1_ (*N*_*K*_ = 1, *N*_*U*_ = 0, *N*_*S*_ = 1). Parentheses show the *Lab–Bayes* result for a *single* stain at the maximum quantity in each case (bold in the Case column).

| Case | Quantities $q_{S1}$ (ng) | $\log_{10} \text{LR}_{S1}$ (Group) | $\log_{10} \text{LR}_{S1}$ (Lab-Bayes) | $\log_{10} \text{LR}_{S1}$ (Lab-Vague) |
| --- | --- | --- | --- | --- |
| 3 stains (all very low) | **0.01**; 0.01; 0.01 | -1.86 | -1.12 (0.61) | -1.71 |
| 3 stains (all low) | 0.04; 0.05; **0.06** | 0.58 | 1.66 (1.68) | 1.66 |
| 3 stains (all medium) | 0.90; 1.00; **1.10** | 3.16 | 4.84 (2.63) | 11.70 |
| 3 stains (all high) | 1.40; 1.50; **1.60** | 3.37 | 5.07 (2.69) | 13.46 |
| 3 stains (all very high) | 4.50; 5.00; **5.50** | 3.69 | 5.47 (2.81) | 19.16 |
| 3 stains (one high, two low) | **1.50**; 0.01; 0.10 | 0.78 | 2.19 (2.68) | 5.05 |
| 3 stains (one very high, two low) | **5.00**; 0.01; 0.10 | 0.76 | 2.32 (2.80) | 6.96 |

*Note.* Replicate stains are treated as conditionally independent given the fitted model parameters.

Mixed scenarios, where one stain yields a high quantity and others are low, produce moderated support. The Lab–Bayes model accumulates evidence without excessive inflation, whereas the Lab–Vague model can produce unrealistically large LRs when multiple high-quantity replicates are observed.

#### 3.3.2. Impact of Specified Unknown Contributors

The presence of specified unknown contributors alters the evidential context for a person of interest in a systematic and predictable way. When unknown contributors are associated with very low DNA quantities, their impact on the likelihood ratio for the person of interest is limited, whereas moderate quantities attributed to the person of interest across one or more stains rapidly dominate. In contrast, unknown contributors associated with high DNA quantities introduce direct competition. Under a single-relevant-actor assumption (*N*_*S*_ = 1), a high-quantity unknown provides a plausible alternative relevant actor (e.g. offender), thereby reducing or reversing support for the person of interest. When multiple contributors exhibit comparable quantities, the likelihood ratio for each approaches unity, reflecting that the evidence supports the proposition that one of the contributors is the relevant actor, but does not discriminate between them.

These effects are demonstrated explicitly in Supplement S5 (Fig. S1), where identical observed quantities are evaluated under different assumptions about the number of relevant actors *N*_*S*_ and unknown contributors

### 3.4. Impact of the Assumed Number of Relevant Actors

Table 4 and Supplement S5 (Fig. S1) both illustrate the critical role of the assumed number of relevant actors, *N*_*S*_. With two contributors exhibiting equally strong evidence, assuming a single relevant actor yields neutral LRs for both. When two relevant actors are assumed, both contributors receive strong support, because the hypothesis that both are relevant actors becomes admissible.

**Table 4:** Effect of number of relevant actors (*N*_*S*_) on log_10_ LR (*Lab–Bayes* Model) for Two Knowns (S1=1.5ng, S2=1.5ng), except for the final row where S2=0.05ng. Single stain simulation.

| Case ID | $N_S$ | $\log_{10}$ LR<sub>S1</sub> | $\log_{10}$ LR<sub>S2</sub> | $\log_{10}$ LR<sub>none</sub> |
| --- | --- | --- | --- | --- |
| S1 and S2 = 1.5ng | 0 | NA | NA | NA |
| S1 and S2 = 1.5ng | 1 | 0.00 | 0.00 | -2.98 |
| S1 and S2 = 1.5ng | 2 | 2.68 | 2.68 | -5.36 |
| S1 = 1.5ng; S2 = 0.05ng | 2 | 2.68 | 1.58 | -4.27 |

This demonstrates that activity-level interpretation is inseparable from contextual assumptions. Changing *N*_*S*_ alters the hypothesis space itself, leading to qualitatively different evidential conclusions even when the observed data remain unchanged.

### 3.5. Implications for Casework

These results show that DNA quantities must be interpreted within a fully specified activity-level context. Laboratory performance, the presence of other contributors, the number of stains, and assumptions about the number of relevant actors all materially affect the resulting likelihood ratios.

The Lab-Bayes approach provides a practical balance between robustness and specificity, avoiding instability while remaining sensitive to laboratory-specific behaviour. The framework discourages selective interpretation in a formal sense: all specified contributors, both known and unknown, are represented explicitly in the exhaustive hypothesis set, and each contributes a likelihood term to the numerator and denominator of the LR. Because the LR is obtained by summing over all admissible elemental hypotheses, it is not possible to condition on a preferred contributor or subset of results without altering the hypothesis space itself.

### 3.6. Impact of Lab-Specific Calibration

To provide practical guidance for laboratories considering adoption of the HaloGen framework, we conducted two sets of simulation studies. Experiment A characterises inter-laboratory variation by comparing pooled and lab-specific models across all available datasets. Experiments B and C then address a pragmatic question: how much in-house data are required for a laboratory to obtain likelihood ratios that are meaningfully more relevant than those produced by a generic pooled model.

### 3.7. Inter-Laboratory Variation in Evidential Weight (Experiment A)

Experiment A compares case-level log_10_ LR values across all 20 datasets from participating laboratories at several DNA quantities. Each data-set was analysed using three modelling strategies: a pooled *Group* model, a lab-specific hierarchical *Lab–Bayes* model, and a standalone *Lab–Vague* model. Results are shown in Fig. 3, with corresponding parameter summaries provided in Supplementary Table S2.

**Figure 3:**
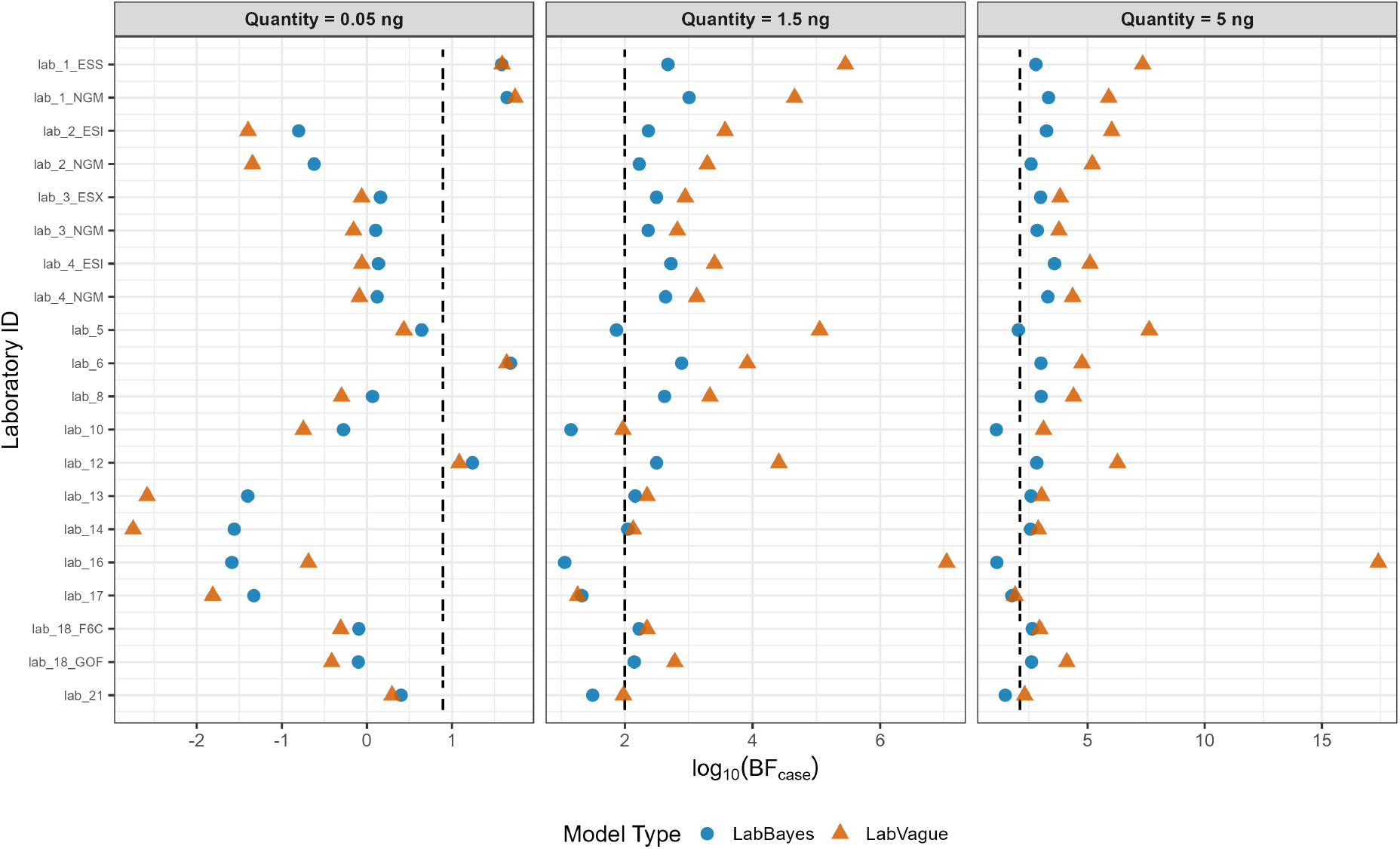
Experiment. **A.** Comparison of log_10_ LR_case_ values across laboratories for multiple DNA quantities. Dashed lines show the pooled *Group* model; blue circles denote *Lab–Bayes* estimates and orange triangles denote *Lab–Vague* estimates. Calculations use the exhaustive *N*_*S*_ = 1 formulation.

Three consistent findings emerge.

First, substantial inter-laboratory variation is observed despite identical experimental protocols. Likelihood ratios differ by several orders of magnitude across laboratories, indicating that transfer efficiency and dropout behaviour vary materially between datasets.

Second, laboratories separate systematically into high- and low-DNA-recovery groups. High-recovery laboratories (e.g. `lab_1_ESS`) tend to produce larger likelihood ratios than the pooled reference, while low-recovery laboratories (e.g. `lab_10`) produce smaller or even negative values. These differences are driven primarily by variation in effective failure rates and the strength of direct transfer.

Third, the choice of modelling strategy matters. The *Lab–Vague* model often yields extreme values at very low and very high quantities, reflecting the absence of cross-laboratory regularisation. The *Lab–Bayes* model moderates these extremes while retaining laboratory-specific behaviour, producing more stable and interpretable results across the full quantity range.

These results motivate the calibration studies that follow, which examine how much laboratory-specific data are required to obtain reliable and context-appropriate likelihood ratios.

### 3.8. Minimum-Effort Calibration for New Laboratories (Experiments B and C)

Experiments B and C investigate how a new laboratory can improve upon the pooled *Group* reference using a limited amount of in-house experimental data. The aim is to quantify how much local data are required before material improvements over the *Group* reference are obtained, under realistic resource constraints. Two laboratories were examined: a high-DNA-recovery laboratory (`lab_1_ESS`) and a low-DNA-recovery laboratory (`lab_10`). Bootstrap analyses (100 replicates) were performed using subsample sizes *n* ∈ {6, 12, 18} at three DNA quantities (0.05, 1.5, and 5 ng), under the exhaustive *N*_*S*_ = 1 framework.

Two modelling configurations were considered:

1. **Experiment B:** *Lab–Bayes* for direct transfer and *Group* for secondary transfer;
2. **Experiment C:** *Lab–Bayes* for both direct and secondary transfer.

The motivation for Experiment B is practical. Direct-transfer experiments are typically simpler to perform for the resource limited laboratory. Experiment B addresses the question of whether incorporating *Lab–Bayes* for direct transfer alone already improves upon exclusive reliance on the *Group* model.

Fig. 4 shows that even a small number of in-house experiments can materially improve evidential relevance relative to exclusive reliance on the *Group* reference. For the low-DNA-recovery laboratory (`lab_10`), the *Group* model systematically overstates evidential weight. Using *Lab–Bayes* for direct transfer with as few as *n* = 6 experiments substantially reduces this inflation and moves likelihood ratios toward the laboratory’s full *Lab–Bayes* benchmark. Introducing *Lab–Bayes* for secondary transfer in Experiment C produces comparatively little additional change for this laboratory, indicating that learning the laboratory’s elevated failure behaviour is the dominant correction.

**Figure 4:**
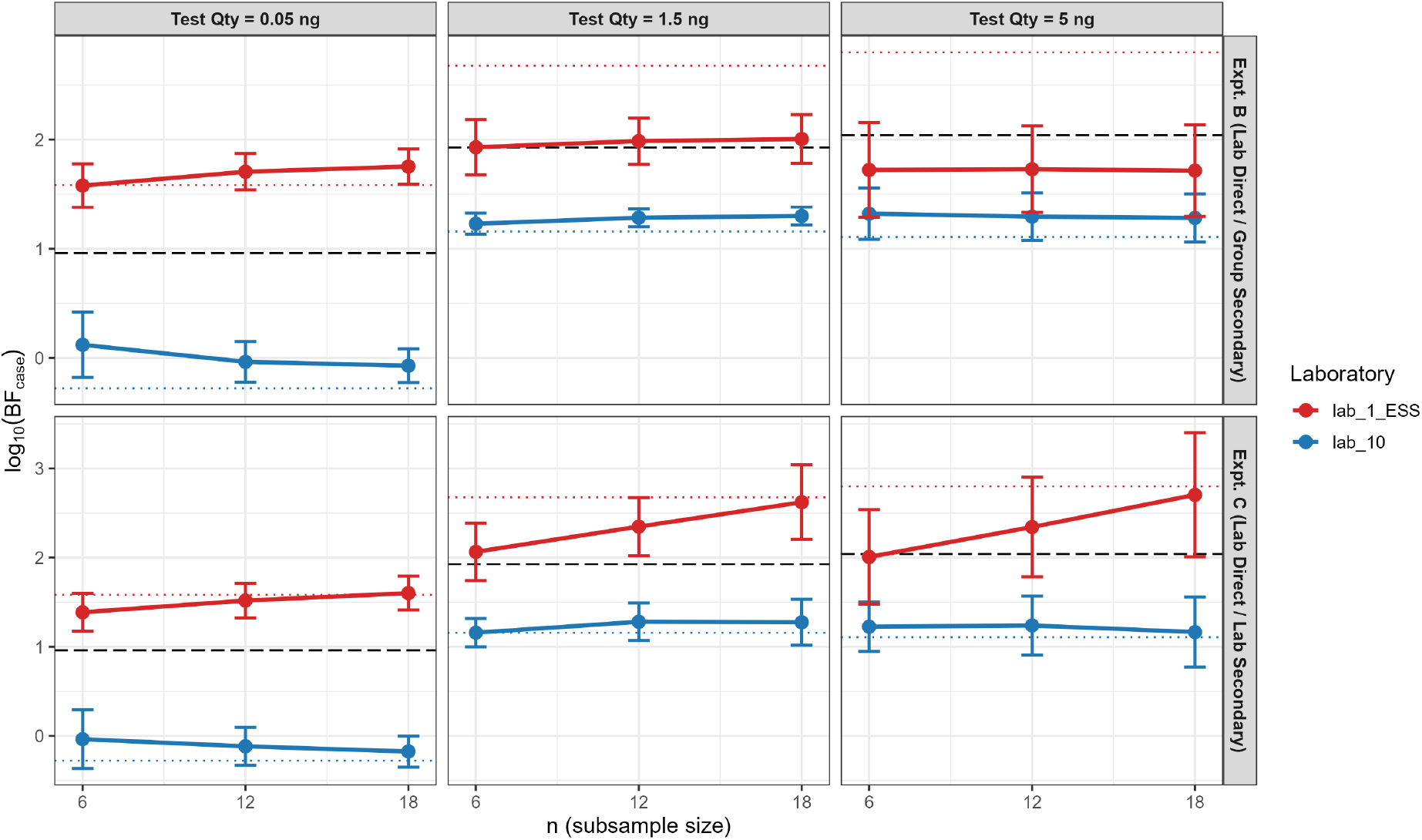
Median ± SD of log_10_(LR_case_) over 100 bootstraps versus subsample size *n*. Rows: Experiment B (top), Experiment C (bottom). Columns: test quantities. Red = `lab_1_ESS`, blue = `lab_10`. Dashed horizontal lines = Group benchmark (lab-agnostic). Dotted horizontal lines = full-data Lab– Bayes benchmark for the corresponding lab. DL = 0.001.

For the high-DNA-recovery laboratory (`lab_1_ESS`), the behaviour differs. When secondary transfer is modelled using *Group* parameters (Experiment B), likelihood ratios are systematically lower than the full *Lab–Bayes* benchmark at moderate and high DNA quantities. Only when Lab–Bayes is used for both direct and secondary transfer (Experiment C) do results converge toward the laboratory’s benchmark. In this setting, using the Group model for secondary transfer leads to conservative estimates because the pooled secondary-transfer distribution assigns too much probability mass to higher quantities than are typical for this laboratory, inflating the denominator of the likelihood ratio.

Across both laboratories, increasing *n* reduces variability and improves alignment with the full *Lab–Bayes* benchmark. In practice, *n* ≈12–18 experiments are often sufficient to obtain substantial gains over the *Group* model. Where resources are limited, replacing *Group* direct transfer with *Lab–Bayes* direct transfer already provides an alternative to relying entirely on pooled cross-laboratory data. Where resources permit, using *Lab– Bayes* for both transfer mechanisms yields the most stable likelihood ratios.

#### Practical implications

These results provide guidance for laboratories wishing to use `HaloGen` with ReAct-like direct/secondary transfer data. They should not be read as universal validation requirements for all activity-level interpretation. External or literature-derived data may be informative where their relevance to the case circumstances is explicitly justified. However, within the `HaloGen` framework, uncritical use of a pooled *Group* model can either overstate or understate evidential strength for a particular laboratory. For laboratories performing below the *Group* reference, limited local calibration of direct transfer can substantially reduce anti-conservative inflation caused by reliance on the pooled model. For laboratories performing above the *Group* reference, calibrating both direct and secondary transfer is more important because the pooled secondary model may otherwise inflate the denominator and produce overly conservative LRs. Where no relevant literature or population data are available, generating a focused laboratory-specific experimental dataset may be more informative than attempting to calibrate against a single external study of uncertain relevance.

Complete numerical results for Experiments B and C are provided in the Supplementary Material (Tables S3 and S4).

## 4. Concluding Remarks

This paper has presented the practical implications of the HaloGen framework for evaluating DNA quantity results under activity-level propositions. At its core, HaloGen combines hierarchical Bayesian modelling with an exhaustive likelihood framework to address a central challenge in forensic science: substantial inter-laboratory variation in DNA transfer, recovery, and dropout.

Analysis of the multi-laboratory ReAct dataset confirms that laboratories differ markedly in transfer efficiency and failure rates, and that these differences translate directly into orders-of-magnitude variation in activity-level likelihood ratios. As a consequence, likelihood ratios derived from pooled or external data can be misleading when applied uncritically to a specific laboratory.

To address this, we examined a series of simulation studies designed to inform practical implementation. Experiment A demonstrated that the generic *Group* model, while useful as a population reference, can either overstate or understate evidential weight depending on a laboratory’s true performance. Experiments B and C showed that modest in-house validation, often as few as 6-18 experiments, can substantially improve relevance and stability. For lower-performing laboratories, learning the direct-transfer behaviour alone is often sufficient to avoid anti-conservative results, whereas higher-performing laboratories benefit from calibrating both direct and secondary transfer, requiring at least *n* = 12.

### 4.1. The Validation Imperative

A central conclusion for quantitative `HaloGen` reporting is that laboratories should not adopt activity-level parameters or likelihood ratios from external sources uncritically. This statement concerns formal model-based LR calculations within the `HaloGen` framework. It does not imply that all activity-level opinions require `HaloGen`-style statistical validation, nor does it preclude expert probability assignments informed by relevant literature, experience, or case-specific judgement. Rather, where a quantitative model is used, the relevance of the calibration data, the activity-level propositions, the sampling design, and the laboratory process must be validated, justified, or explored through sensitivity analysis.

The model also does not eliminate scientific judgement. The expert must still decide whether the available transfer, prevalence, persistence, and recovery (TPPR) data are relevant to the case circumstances, whether external data are suitable proxies, which propositions should be compared, and how uncertainty in sampling, transfer opportunity, persistence, and recovery should be communicated. `HaloGen` makes these assumptions explicit and quantifies their consequences; it does not replace the expert’s responsibility to justify them.

Even small mismatches between laboratories can lead to systematic overstatement or understatement of evidential strength. `HaloGen`’s hierarchical structure mitigates this risk by combining limited local data with broader population information, allowing likelihood ratios to adjust in a data-driven manner while explicitly reflecting remaining uncertainty in laboratory-specific performance.

### 4.2. Implications for Casework and Confirmation Bias

The `HaloGen` framework has important implications for casework practice. By construction, it requires all contributors, known and unknown, and all stains, including non-detects, to be evaluated jointly given clearly specified activity propositions. This structure reduces the risk of confirmation bias, where attention is focused on results supporting a favoured individual while background or conflicting findings are discounted. The Birgitte Tengs case [8] illustrates this risk. Although DNA recovered from two samples aligned with the suspect, the majority of samples were either uninformative or contained DNA assigned to unknown contributors. Such findings should not be treated as automatically neutral; their effect depends on the propositions, the sampling strategy, and the full evidence set. A framework such as `HaloGen` makes the consequences of these assumptions explicit by requiring all specified contributors and non-detections to be considered within the hypothesis space.

The importance of context is further illustrated by cases involving the assumed number of relevant actors. When *N*_*S*_ = 0, any detected DNA contradicts the proposition, yielding a likelihood ratio of zero for individual involvement. This is not merely theoretical, as shown by the miscarriage of justice in the Farah Jamah case [9], where DNA was detected despite no offence having occurred. Conversely, in cases such as Amanda Knox and Raffaele Sollecito [10], competing assumptions about *N*_*S*_ lead to fundamentally different interpretations. HaloGen allows such assumptions to be modelled explicitly, making their impact transparent to the court.

### 4.3. Future Directions

HaloGen is designed as a modular and extensible framework. Future work will focus on incorporating additional evidence types (e.g. body-fluid tests, time since deposition), improving estimation of failure rates using casework data, and developing priors for background DNA based on real-world sampling rather than sterile experimental conditions. Greater standardisation of laboratory practices would further strengthen the population-level components of the model.

### 4.4. Final Remarks

HaloGen provides a coherent, transparent, and empirically grounded framework for interpreting DNA quantity results given activity-level propositions. By integrating multiple stains, contributors, and contextual assumptions within a single likelihood ratio, it moves beyond *ad hoc* approaches and offers a defensible basis for forensic inference. The framework does not remove the need for judgement, but it makes the consequences of that judgement explicit, which is an essential requirement for fair and balanced evaluation of DNA findings given activity-level propositions in court.

## Acknowledgements

Funding for this Project was received from the European Union’s Internal Security Fund -Police (ISFP) – with Grant Agreement title: Competency, Education, Research, Testing, Accreditation, and Innovation in Forensic Science” [CERTAIN-FORS] ISFP-2020-AG-IBA-ENFSI and number: 101051099. Also: ReAct II – Extension of ReAct. Work package 5 within the FOR-FUTURE project - Forensic Fundamentals, Technology, Multidisciplinarity, Research, Evaluation (FOR-FUTURE) — ISF-2023-TF2-AG-ENFSI-IBA-2.

