## Supplementary Files for "Likelihood Ratios Given Activity-Level Propositions for DNA Transfer Evidence: Practical Implementation and Simulation Studies Using the HaloGen Engine (Part II)"

### S1. Notation and symbols

Table S1: Notation and Symbols

| Symbol | Type | Meaning / usage |
|---|---|---|
| | evidence | Observed case evidence used in the LR calculation, including contributor-level DNA quantities and any modelled non-detect information relevant to the propositions. |
| $q$ | scalar | Observed DNA quantity (ng) for a single contributor. For a detected contributor quantity, $q \geq \mathrm{DL}$; values $q < \mathrm{DL}$ are treated as non-detects. |
| DL | scalar | Detection limit used in the activity-level quantity model. |
| $(\mu, \sigma, k)$ | parameters | Parameters of the lab- or group-specific zero-augmented lognormal transfer model: mean $\mu$, standard deviation $\sigma$, and zero-transfer/no-transfer probability $k$. |
| $F_0$ | parameter | Case-level non-detection probability for a relevant but unobserved actor; used when an elemental hypothesis requires an actor who left no detectable DNA. |
| $N_S$ | integer | Number of relevant actors posited under a given activity-level proposition. |
| $N_K$, $N_U$ | integers | Numbers of specified known contributors and specified unknown contributors. |
| $S_i$, $U_j$ | indices | Specified known contributor $S_i$ and specified unknown contributor $U_j$, respectively. |
| $q_{S_i}$, $q_{U_j}$ | scalar or vector | Measured quantities (ng) attributed to the corresponding contributor across one or more stains. |
| $M_x$ | scalar | Mixture proportion of contributor $x$ within a stain, where relevant to estimating contributor-level quantities. |
| $t(q)(q)$, $s(q)(q)$ | functions | Per-draw case-level likelihood contributions for a detected quantity $q \geq \mathrm{DL}$: $t(q)(q)$ for the direct-transfer pathway and $s(q)(q)$ for the secondary-transfer pathway. For detected quantities, both are evaluated using conditional-on-detection densities. Non-detects are handled separately through probability-mass terms, including $F_0$ where an elemental hypothesis requires an unobserved actor, and are not treated as zero-valued densities. |
| LR | ratio | Likelihood ratio; in this paper equal to the Bayes factor under equal elemental priors. |
| $\mathrm{LR}_{\mathrm{case}}$ | ratio | Case-level LR computed by exhaustive summation over all admissible elemental hypotheses for fixed $N_S$. |
| $\mathrm{LR}_{\mathrm{case},X}$ | ratio | Case-level LR for contributor $X$ (e.g., $S_1$, $U_3$) obtained by partitioning elemental hypotheses according to whether $X$ is assigned to the direct-transfer actor role. |
| $\log_{10}\mathrm{LR}_{\mathrm{case},X}$ | scalar | $\log_{10}$-transformed case-level LR for contributor $X$, typically reported as the evidential weight. |
| *Group*, *Lab–Bayes*, *Lab–Vague* | labels | Model types: pooled lab-agnostic, hierarchical lab-specific, and standalone lab-specific, respectively. |

### S2. Model parameters calculated for all labs

Table S2: Comparison of $\log_{10}$LR values for the Group, Lab-Bayes, and Lab-Vague models across all 20 laboratories for five different single-stain quantities ($q$).

| **Lab ID** | **Model** | $q = 0.01$ | $q = 0.05$ | $q = 1.0$ | $q = 1.5$ | $q = 5.0$ |
|---|---|---|---|---|---|---|
| lab_1_ESS | Group | 0.09 | 0.89 | 1.92 | 2.00 | 2.12 |
| | LabBayes | 0.61 | 1.58 | 2.61 | 2.68 | 2.80 |
| | LabVague | 0.42 | 1.59 | 4.89 | 5.47 | 7.37 |
| lab_1_NGM | Group | 0.09 | 0.89 | 1.92 | 2.00 | 2.12 |
| | LabBayes | 0.69 | 1.64 | 2.88 | 3.01 | 3.34 |
| | LabVague | 0.59 | 1.74 | 4.26 | 4.65 | 5.90 |
| lab_2_ESI | Group | 0.09 | 0.89 | 1.92 | 2.00 | 2.12 |
| | LabBayes | −2.56 | −0.81 | 2.05 | 2.37 | 3.26 |
| | LabVague | −2.69 | −1.39 | 2.83 | 3.57 | 6.02 |
| lab_2_NGM | Group | 0.09 | 0.89 | 1.92 | 2.00 | 2.12 |
| | LabBayes | −2.82 | −0.63 | 2.03 | 2.23 | 2.59 |
| | LabVague | −3.10 | −1.35 | 2.69 | 3.29 | 5.19 |
| lab_3_ESX | Group | 0.09 | 0.89 | 1.92 | 2.00 | 2.12 |
| | LabBayes | −1.38 | 0.16 | 2.29 | 2.50 | 2.99 |
| | LabVague | −1.71 | −0.06 | 2.63 | 2.94 | 3.82 |
| lab_3_NGM | Group | 0.09 | 0.89 | 1.92 | 2.00 | 2.12 |
| | LabBayes | −1.36 | 0.10 | 2.17 | 2.37 | 2.85 |
| | LabVague | −1.65 | −0.16 | 2.49 | 2.82 | 3.78 |
| lab_4_ESI | Group | 0.09 | 0.89 | 1.92 | 2.00 | 2.12 |
| | LabBayes | −1.09 | 0.13 | 2.43 | 2.72 | 3.60 |
| | LabVague | −0.97 | −0.06 | 2.89 | 3.40 | 5.10 |
| lab_4_NGM | Group | 0.09 | 0.89 | 1.92 | 2.00 | 2.12 |
| | LabBayes | −1.32 | 0.12 | 2.40 | 2.64 | 3.31 |
| | LabVague | −1.41 | −0.09 | 2.72 | 3.12 | 4.36 |
| lab_5 | Group | 0.09 | 0.89 | 1.92 | 2.00 | 2.12 |
| | LabBayes | −0.30 | 0.64 | 1.78 | 1.87 | 2.05 |
| | LabVague | −0.44 | 0.43 | 4.29 | 5.06 | 7.63 |
| lab_6 | Group | 0.09 | 0.89 | 1.92 | 2.00 | 2.12 |
| | LabBayes | 0.62 | 1.68 | 2.81 | 2.88 | 3.02 |
| | LabVague | 0.52 | 1.64 | 3.64 | 3.92 | 4.77 |
| lab_8 | Group | 0.09 | 0.89 | 1.92 | 2.00 | 2.12 |
| | LabBayes | −1.80 | 0.06 | 2.43 | 2.62 | 3.02 |
| | LabVague | −2.29 | −0.30 | 2.95 | 3.33 | 4.40 |
| lab_10 | Group | 0.09 | 0.89 | 1.92 | 2.00 | 2.12 |
| | LabBayes | −1.77 | −0.28 | 1.11 | 1.16 | 1.11 |
| | LabVague | −1.71 | −0.75 | 1.61 | 1.97 | 3.11 |

*Continued on next page*

Table S2: (continued)

| **Lab ID** | **Model** | $q = 0.01$ | $q = 0.05$ | $q = 1.0$ | $q = 1.5$ | $q = 5.0$ |
|---|---|---|---|---|---|---|
| lab_12 | Group | 0.09 | 0.89 | 1.92 | 2.00 | 2.12 |
| | LabBayes | 0.46 | 1.23 | 2.38 | 2.50 | 2.83 |
| | LabVague | 0.43 | 1.08 | 3.86 | 4.41 | 6.28 |
| lab_13 | Group | 0.09 | 0.89 | 1.92 | 2.00 | 2.12 |
| | LabBayes | −4.20 | −1.41 | 1.92 | 2.16 | 2.59 |
| | LabVague | −6.35 | −2.58 | 2.00 | 2.35 | 3.04 |
| lab_14 | Group | 0.09 | 0.89 | 1.92 | 2.00 | 2.12 |
| | LabBayes | −4.31 | −1.57 | 1.77 | 2.04 | 2.56 |
| | LabVague | −6.35 | −2.75 | 1.76 | 2.13 | 2.90 |
| lab_16 | Group | 0.09 | 0.89 | 1.92 | 2.00 | 2.12 |
| | LabBayes | −4.07 | −1.59 | 0.92 | 1.06 | 1.13 |
| | LabVague | 6.36 | −0.69 | 4.43 | 7.04 | 17.42 |
| lab_17 | Group | 0.09 | 0.89 | 1.92 | 2.00 | 2.12 |
| | LabBayes | −3.28 | −1.33 | 1.11 | 1.32 | 1.78 |
| | LabVague | −3.85 | −1.81 | 0.98 | 1.26 | 1.91 |
| lab_18_F6C | Group | 0.09 | 0.89 | 1.92 | 2.00 | 2.12 |
| | LabBayes | −1.73 | −0.09 | 2.03 | 2.22 | 2.64 |
| | LabVague | −1.93 | −0.30 | 2.11 | 2.34 | 2.95 |
| lab_18_GOF | Group | 0.09 | 0.89 | 1.92 | 2.00 | 2.12 |
| | LabBayes | −1.56 | −0.10 | 1.95 | 2.14 | 2.61 |
| | LabVague | −1.63 | −0.41 | 2.37 | 2.79 | 4.13 |
| lab_21 | Group | 0.09 | 0.89 | 1.92 | 2.00 | 2.12 |
| | LabBayes | −0.70 | 0.40 | 1.45 | 1.50 | 1.49 |
| | LabVague | −0.85 | 0.30 | 1.83 | 1.97 | 2.31 |

### S3. Parameters calculated for Experiments B and C

Table S3: Experiment B (Lab Direct / Group Secondary): Median and SD of $\log_{10}(\mathrm{LR}_{\mathrm{case}})$ over 100 bootstraps, compared with Group and Lab–Bayes benchmarks and the lab-specific fail rate $F_{0,\mathrm{lab}}$.

| **Lab ID** | **Qty (ng)** | $\boldsymbol{F_{0,\mathrm{lab}}}$ | **Benchmark** $\log_{10}$ LR | | **Partial Lab Model** ($n = 6$) | **Partial Lab Model** ($n = 12$) | **Partial Lab Model** ($n = 18$) |
|---|---|---|---|---|---|---|---|
| | | | **Group** | **Lab–Bayes** | **Median** $\pm$ **SD** | **Median** $\pm$ **SD** | **Median** $\pm$ **SD** |
| **Lab: lab_1_ESS** | | | | | | | |
| | 0.05 | 0.022 | 0.82 | 1.58 | $1.58 \pm 0.18$ | $1.69 \pm 0.17$ | $1.74 \pm 0.17$ |
| | 1.50 | 0.022 | 1.86 | 2.68 | $1.95 \pm 0.21$ | $2.00 \pm 0.21$ | $2.04 \pm 0.19$ |
| | 5.00 | 0.022 | 1.96 | 2.80 | $1.73 \pm 0.37$ | $1.79 \pm 0.37$ | $1.78 \pm 0.33$ |
| **Lab: lab_10** | | | | | | | |
| | 0.05 | 0.420 | 0.82 | $-0.28$ | $0.11 \pm 0.24$ | $-0.02 \pm 0.17$ | $-0.07 \pm 0.18$ |
| | 1.50 | 0.420 | 1.86 | 1.16 | $1.23 \pm 0.08$ | $1.30 \pm 0.07$ | $1.30 \pm 0.08$ |
| | 5.00 | 0.420 | 1.96 | 1.10 | $1.31 \pm 0.17$ | $1.30 \pm 0.18$ | $1.31 \pm 0.22$ |

Table S4: Experiment C (Lab Direct / Lab Secondary): Median and SD of $\log_{10}(\mathrm{LR}_{\mathrm{case}})$ over 100 bootstraps, compared with Group and Lab–Bayes benchmarks and the lab-specific fail rate $F_{0,\mathrm{lab}}$.

| **Lab ID** | **Qty (ng)** | $\boldsymbol{F_{0,\mathrm{lab}}}$ | **Benchmark** $\log_{10}$ LR | | **Partial Lab Model** ($n = 6$) | **Partial Lab Model** ($n = 12$) | **Partial Lab Model** ($n = 18$) |
|---|---|---|---|---|---|---|---|
| | | | **Group** | **Lab–Bayes** | **Median** $\pm$ **SD** | **Median** $\pm$ **SD** | **Median** $\pm$ **SD** |
| **Lab: lab_1_ESS** | | | | | | | |
| | 0.05 | 0.022 | 0.82 | 1.58 | $1.40 \pm 0.18$ | $1.51 \pm 0.20$ | $1.57 \pm 0.19$ |
| | 1.50 | 0.022 | 1.86 | 2.68 | $2.08 \pm 0.29$ | $2.28 \pm 0.35$ | $2.59 \pm 0.38$ |
| | 5.00 | 0.022 | 1.96 | 2.80 | $2.04 \pm 0.48$ | $2.30 \pm 0.57$ | $2.71 \pm 0.61$ |
| **Lab: lab_10** | | | | | | | |
| | 0.05 | 0.420 | 0.82 | $-0.28$ | $-0.01 \pm 0.27$ | $-0.14 \pm 0.20$ | $-0.17 \pm 0.19$ |
| | 1.50 | 0.420 | 1.86 | 1.16 | $1.22 \pm 0.15$ | $1.24 \pm 0.21$ | $1.28 \pm 0.22$ |
| | 5.00 | 0.420 | 1.96 | 1.10 | $1.31 \pm 0.24$ | $1.21 \pm 0.30$ | $1.22 \pm 0.36$ |

### S4. A validation of priors

#### *S4.1. Choice of Priors for Bayesian Models*

HaloGen employs weakly informative priors designed to stabilise inference without overwhelming the experimental data. We distinguish clearly between (i) the *within-laboratory* non-detect probability $k$ used in the transfer likelihood and (ii) the *case-level* fail-rate parameter $F_0$ used in the hypothesis set. This separation is essential for coherent activity-level inference. The priors used for the Group, Lab–Bayes, and Lab–Vague models are summarised below; the $F_0$ policy is applied uniformly across all models.

*Transfer likelihood: zero-augmented model with censoring.* For a given laboratory and transfer mechanism, direct or secondary, the experimental DNA quantity $q$ is modelled using a zero-augmented lognormal mixture with parameters $\theta = (\mu, \sigma, k)$ and detection limit DL. A recorded non-detect can arise either from a zero-transfer/no-transfer event, with probability $k$, or from a positive transfer that falls below the detection limit. The probability mass assigned to a non-detect is therefore

$$P(q < \mathrm{DL} \mid \theta, \mathrm{DL}) = k + (1-k)\,\Phi\left(\frac{\log \mathrm{DL} - \mu}{\sigma}\right),$$

where $\Phi(\cdot)$ is the standard normal distribution function. Equivalently, if $F(\mathrm{DL})$ denotes the lognormal distribution function evaluated at the detection limit, this can be written as

$$P(q < \mathrm{DL} \mid \theta, \mathrm{DL}) = k + (1-k)F(\mathrm{DL}).$$

For an observed detected experimental quantity $q_{\mathrm{obs}} \geq \mathrm{DL}$, the likelihood contribution used when fitting the transfer model is the continuous density

$$L_{\mathrm{fit}}(q_{\mathrm{obs}} \mid \theta, \mathrm{DL}) = (1-k)\, f_{\mathrm{LN}}(q_{\mathrm{obs}} \mid \mu, \sigma), \qquad q_{\mathrm{obs}} \geq \mathrm{DL},$$

where $f_{\mathrm{LN}}$ denotes the lognormal density. Thus, in the experimental fitting likelihood, non-detects contribute through a probability mass, whereas observed detected quantities contribute through a density.

This experimental fitting likelihood should be distinguished from the case-level likelihood used to evaluate a detected contributor quantity. At case level, once a contributor quantity $q \geq \mathrm{DL}$ has been observed, HaloGen uses the conditional-on-detection density defined in Part I [1], Section 2.3:

$$f_{\mathrm{cond}}(q) = \frac{f_{\mathrm{LN}}(q \mid \mu, \sigma)}{1 - F(\mathrm{DL})} = \frac{f_{\mathrm{LN}}(q \mid \mu, \sigma)}{1 - \Phi((\log \mathrm{DL} - \mu)/\sigma)}, \qquad q \geq \mathrm{DL}.$$

Here $F(\mathrm{DL}) = \Phi((\log \mathrm{DL} - \mu)/\sigma)$. The direct and secondary case-level components are then

$$t_c(q) = f_{D,\mathrm{cond}}(q), \qquad s_c(q) = f_{S,\mathrm{cond}}(q), \qquad q \geq \mathrm{DL}.$$

Thus, the zero-augmented mixture likelihood is used to estimate the experimental transfer parameters, whereas detected case quantities are evaluated using conditional-on-detection densities.

For each Monte Carlo index $m$, HaloGen uses posterior parameter draws for the direct and secondary transfer arms,

$$\theta_D^{(m)} = (\mu_D^{(m)}, \sigma_D^{(m)}, k_D^{(m)}), \qquad \theta_S^{(m)} = (\mu_S^{(m)}, \sigma_S^{(m)}, k_S^{(m)}),$$

and evaluates the corresponding case-level likelihood components $t_c^{(m)}(q)$ and $s_c^{(m)}(q)$ using the same draw index. This paired per-draw construction propagates uncertainty in both transfer arms coherently into the posterior Monte Carlo distribution of the LR/BF.

Non-detects are handled separately through probability-mass terms, including the actor non-detection probability $F_0$ where an elemental hypothesis requires a relevant actor who is not represented among the detected contributors. This distinction between the experimental transfer likelihood and the case-level likelihood is developed formally in Part I [1], Sections 2.3–2.4.

Case-level likelihood ratios are computed from these paired posterior draws and reported as medians with 10–90% intervals on the $\log_{10}$ scale.

*Group model (hierarchical priors).* Laboratory-specific parameters are modelled hierarchically using non-centred parameterisations:

$$\mu_{\text{lab}} = \mu_0 + \tau_\mu \, \mu^*_{\text{lab}}, \qquad \mu^*_{\text{lab}} \sim \mathcal{N}(0,1),$$

with analogous forms for $\log \sigma_{\text{lab}}$. Hyperpriors are specified as

$$\mu_0 \sim \mathcal{N}(0, 10^2), \qquad \tau_\mu \sim \text{Student-}t_3(0,5),$$
$$\log \sigma_0 \sim \mathcal{N}(0, 10^2), \qquad \tau_{\log \sigma} \sim \text{Student-}t_3(0,5).$$

The dropout probability is modelled via a Beta mean–concentration hierarchy:

$$k_{\text{lab}} \mid \mu_k, \phi_k \sim \text{Beta}(\mu_k \phi_k, (1-\mu_k)\phi_k),$$

with

$$\mu_k \sim \text{Beta}(2,2), \qquad \phi_k \sim \text{Student-}t_3(0,5).$$

This structure provides partial pooling across laboratories while allowing substantial heterogeneity in transfer behaviour.

*Lab–Bayes model.* The Lab–Bayes model inherits the hierarchical structure of the Group model. Posterior summaries of the Group hyperparameters $(\mu_0, \tau_\mu, \log \sigma_0, \tau_{\log \sigma}, \mu_k, \phi_k)$ are used as fixed inputs when fitting the laboratory-specific model. This implements partial pooling: each laboratory's inference is informed by population-level behaviour while remaining responsive to its own data.

*Lab–Vague model.* The Lab–Vague model is fitted independently for each laboratory using weak, non-hierarchical priors implemented via the same non-centred parameterisation:

$$\mu_0 = 0, \quad \tau_\mu = 10; \qquad \log \sigma_0 = 0, \quad \tau_{\log \sigma} = 5.$$

For the dropout probability,

$$\mu_k = 0.5, \quad \phi_k = 1,$$

which implies a $\text{Beta}(0.5, 0.5)$ prior for $k_{\text{lab}}$. This prior is intentionally broad and places substantial mass near 0 and 1, reflecting the absence of cross-laboratory regularisation in this model. As shown in the Results section, this can lead to instability at the extremes when laboratory datasets are small.

*Case-level fail-rate parameter $F_0$.* The probability that a relevant unobserved actor leaves no detectable DNA is treated separately from the within-lab dropout probability $k$. A Jeffreys-type prior is used:

$$F_0 \sim \text{Beta}(0.5, 0.5),$$

with Monte Carlo draws constrained to a conservative interval $[J, 1 - J]$, where $J$ is determined empirically from direct-transfer data. This policy prevents unrealistically small values of $F_0$ from producing anti-conservative likelihood ratios while preserving inter-laboratory variability.

*Notes on implementation.* All posterior inference is performed using MCMC sampling in the transfer-level model domain. That is, the MCMC procedure estimates posterior distributions for the direct- and secondary-transfer parameters $\theta_D = (\mu_D, \sigma_D, k_D)$ and $\theta_S = (\mu_S, \sigma_S, k_S)$.

In the case-level domain, likelihood ratios are then computed per posterior draw using paired direct- and secondary-transfer parameter draws. For draw $m$, HaloGen evaluates the case-level likelihood components using $\theta_D^{(m)}$ and $\theta_S^{(m)}$, and forms the corresponding LR/BF draw. This paired per-draw construction propagates transfer-parameter uncertainty coherently into the case-level LR distribution.

Summary statistics, namely the median and 10–90% intervals, are reported for presentation only and do not constitute a separate inferential procedure.

*Notes on choices.* (i) We use half-Student-$t$ priors for hierarchical scale parameters to avoid the instability associated with heavy-tailed half-Cauchy distributions. In practice, $\nu = 3$ with scale 5 provides sufficient regularisation while remaining weakly informative across laboratories.

(ii) For the Lab–Vague model, the non-detect probability $k$ is assigned a $\text{Beta}(0.5, 0.5)$ prior, reflecting minimal prior information and allowing the data to dominate. This deliberately exposes the sensitivity of standalone fits to limited sample sizes and contrasts with the stabilising effect of hierarchical shrinkage in the Group and Lab–Bayes models.

(iii) The case-level fail-rate parameter $F_0$ is conceptually and mathematically distinct from the within-laboratory dropout probability $k$. It is assigned a Jeffreys-type $\text{Beta}(0.5, 0.5)$ prior (with clamping) and should not be replaced by optimistic low-$F_0$ priors, which would risk anti-conservative likelihood ratios.

*Sensitivity and reporting.* Likelihood ratios are computed using paired posterior draws and reported as medians with 10–90% credible intervals on the $\log_{10}$ scale. Additional sensitivity analyses exploring alternative priors for $k$ (e.g. $\text{Beta}(1, 1)$ or $\text{Beta}(3, 3)$) yield similar median LRs, with interval width reflecting prior concentration. A limitations note is flagged if the $\log_{10} \text{LR}_{\text{none}}$ interval exceeds three units or if the effective number of paired draws is low.

*Detection-limit robustness*

The censored-mixture likelihood used in HaloGen depends explicitly on the analytical detection limit (DL) through the probability mass assigned to non-detections. To assess the numerical robustness of likelihood ratios to moderate misspecification of the DL, we conducted a dedicated *DL robustness* validation (Section B8 of the `Validation.R` script).

For each diagnostic scenario, likelihood ratios were recomputed under two perturbed detection limits,

$$\text{DL}_{\text{low}} = \tfrac{1}{2}\text{DL} \quad \text{and} \quad \text{DL}_{\text{high}} = 2\text{DL},$$

Table S5: Priors used in HaloGen (as implemented in `Gemini_v14_v6`). Parameters are per lab and transfer mode unless labelled as hyperparameters.

| Model | Parameter | Prior | Rationale / Notes |
|---|---|---|---|
| ***Group (hierarchical / partial pooling across labs)*** | | | |
| | $\mu_{\text{lab}}$ | $\mu_{\text{lab}} = \mu_0 + \tau_\mu \mu^*_{\text{lab}}$, $\mu^*_{\text{lab}} \sim \mathcal{N}(0,1)$, $\mu_0 \sim \mathcal{N}(s_\mu, 10^2)$, $\tau_\mu \sim \text{Student-}t_3(0,5)$ | Weakly informative hierarchical location prior. $s_\mu$ is a fixed offset by transfer type (direct: 0; secondary: $-2.2$). |
| | $\log \sigma_{\text{lab}}$ | $\log \sigma_{\text{lab}} = \log \sigma_0 + \tau_{\log \sigma} \sigma^*_{\text{lab}}$, $\sigma^*_{\text{lab}} \sim \mathcal{N}(0,1)$, $\log \sigma_0 \sim \mathcal{N}(s_\sigma, 10^2)$, $\tau_{\log \sigma} \sim \text{Student-}t_3(0,5)$ | Log-scale ensures $\sigma > 0$. $s_\sigma$ fixed by transfer type (direct: 0; secondary: 0.5). |
| | $k_{\text{lab}}$ | $k_{\text{lab}} \sim \text{Beta}(\mu_k \phi_k, (1-\mu_k)\phi_k)$, $\mu_k \sim \text{Beta}(2,2)$, $\phi_k \sim \text{Student-}t_3(0,5)$ | Mean–concentration hierarchy for dropout. Partial pooling stabilises $k$ across labs. |
| ***Lab–Bayes (lab-specific, inherits Group structure)*** | | | |
| | $\mu_{\text{lab}}, \log \sigma_{\text{lab}}, k_{\text{lab}}$ | Same hierarchical forms as Group, with hyperparameters fixed to Group posterior summaries. | Implements partial pooling ("borrowing strength") from the population while remaining lab-specific. |
| ***Lab–Vague (standalone, weakly constrained)*** | | | |
| | $\mu_{\text{lab}}$ | $\mu_{\text{lab}} = \mu_0 + \tau_\mu \mu^*_{\text{lab}}$, $\mu^*_{\text{lab}} \sim \mathcal{N}(0,1)$, $\mu_0 = 0, \quad \tau_\mu = 10$ | Broad location prior on log-quantity scale. |
| | $\log \sigma_{\text{lab}}$ | $\log \sigma_0 = 0, \quad \tau_{\log \sigma} = 5$ | Allows wide dispersion without heavy tails. |
| | $k_{\text{lab}}$ | $\text{Beta}(0.5, 0.5)$ | Deliberately weak prior; places mass near 0 and 1. Leads to tail inflation when $n$ is small. |
| ***Case level (hypothesis set, shared across models)*** | | | |
| | $F_0$ | $\text{Beta}(0.5, 0.5)$, clamped to $[J, 1-J]$ | Jeffreys-type prior; distinct from $k$. Prevents anti-conservative extremes in case-level LRs. |

and compared with the reference analysis using the nominal DL. Robustness was assessed by examining the absolute difference in the resulting $\log_{10}(\mathrm{LR})$ values between the $\mathrm{DL}_{\mathrm{low}}$ and $\mathrm{DL}_{\mathrm{high}}$ runs.

Three diagnostic scenarios were considered:

1. **Base:** a representative mixture of positive quantities and non-detections, reflecting typical experimental data.
2. **Zeros-only:** all observations set to zero (non-detects), stressing the censoring and dropout component of the likelihood.
3. **Positive-only:** all observations above DL, stressing the continuous lognormal component.

For each scenario, likelihood ratios were computed using paired posterior draws, and the absolute difference

$$|\log_{10} \mathrm{LR}(\mathrm{DL}_{\mathrm{high}}) - \log_{10} \mathrm{LR}(\mathrm{DL}_{\mathrm{low}})|$$

was used as the robustness metric. Small differences indicate that the evidential conclusions are stable to plausible uncertainty in the analytical detection limit, while larger differences flag scenarios in which DL misspecification could materially affect interpretation.

In the base and positive-only scenarios, medians and bands were invariant within numerical tolerance; in the zeros-only scenario, small shifts were observed but remained within predefined acceptance thresholds (Table S7). All rows passed the B8 validation criteria.

Table S6: Detection-limit robustness (Validation B8). Medians and 10–90% bands of the signed paired per-draw difference in $\log_{10}(\mathrm{BF}_{\mathrm{case}})$ between analyses using $\mathrm{DL}_{\mathrm{low}} = \mathrm{DL}/2$ and $\mathrm{DL}_{\mathrm{high}} = 2\mathrm{DL}$, defined as $\Delta_{\mathrm{DL}} = \log_{10} \mathrm{BF}_{\mathrm{case}}(\mathrm{DL}_{\mathrm{high}}) - \log_{10} \mathrm{BF}_{\mathrm{case}}(\mathrm{DL}_{\mathrm{low}})$. Results are shown for three diagnostic scenarios and each model.

| Scenario | Model | Median | $q_{10}$ | $q_{90}$ |
|---|---|---|---|---|
| base | GROUP | −0.019 | −1.000 | 0.037 |
| base | LAB–BAYES | 0.010 | −0.001 | 0.019 |
| base | LAB–VAGUE | 0.031 | 0.011 | 0.055 |
| zeros-only | GROUP | −0.226 | −0.567 | −0.103 |
| zeros-only | LAB–BAYES | −0.208 | −0.894 | −0.106 |
| zeros-only | LAB–VAGUE | −0.285 | −1.340 | −0.115 |
| positive-only | GROUP | −0.019 | −1.000 | 0.037 |
| positive-only | LAB–BAYES | 0.010 | −0.001 | 0.019 |
| positive-only | LAB–VAGUE | 0.031 | 0.011 | 0.055 |

*Implementation note.* For reproducibility, B8 should be run with the same posterior draws and paired per-draw engine as the main analyses, changing only DL. We recommend fixing the random seed and reporting the exact DL values used, together with the model type (Group, Lab–Bayes, Lab–Vague) and the $F_0$ policy (empirical Beta$(0.5, 0.5)$ + clamp).

Table S7: B8 acceptance thresholds for the absolute values of the signed detection-limit differences in Table S6. The thresholds are applied to $|\Delta_{\mathrm{DL,median}}|$, $|\Delta_{\mathrm{DL},q10}|$, and $|\Delta_{\mathrm{DL},q90}|$ for paired per-draw $\log_{10} \mathrm{BF}_{\mathrm{case}}$ summaries. All results passed.

| Bucket | $\lvert\Delta_{\mathrm{DL,median}}\rvert$ | $\lvert\Delta_{\mathrm{DL},q10}\rvert$ | $\lvert\Delta_{\mathrm{DL},q90}\rvert$ |
|---|---|---|---|
| Base, positive-only (all models) | $\leq 0.005$ | $\leq 0.010$ | $\leq 0.010$ |
| Zeros-only (*Group*) | $\leq 0.050$ | $\leq 0.200$ | $\leq 0.200$ |
| Zeros-only (*Lab–Bayes* / *Lab–Vague*) | $\leq 0.150$ | $\leq 0.200$ | $\leq 0.200$ |

### S5. Sensitivity of the Likelihood Ratio to DNA Quantity and Case Context

This supplement provides an illustrative sensitivity analysis of how the case-level likelihood ratio (LR) for a person of interest (S1) depends jointly on DNA quantity, mixture composition, the number of specified unknown contributors, and the assumed number of relevant actors. The purpose is to visualise, in a single setting, the contextual dependencies that underlie the results reported in the main text.

We conducted 1,000 Monte Carlo simulations for a single-stain scenario using the high-DNA-recovery laboratory `lab_1_ESS`. For each simulation, DNA quantities were generated under the HaloGen model and the case-level LR for S1 was computed using paired posterior draws. Points in Fig. S1 show the median $\log_{10}$ LR for S1 as a function of the realised quantity $q_{S1}$.

Columns in the figure vary the number of specified unknown contributors ($N_U$) and the assumed number of relevant actors ($N_S$). Rows compare the three modelling approaches used throughout the paper: the pooled *Group* model, the hierarchical *Lab–Bayes* model, and the standalone *Lab–Vague* model. Point colour indicates the realised mixture proportion of S1 ($M_x$), reflecting the share of the total DNA attributed to the known contributor.

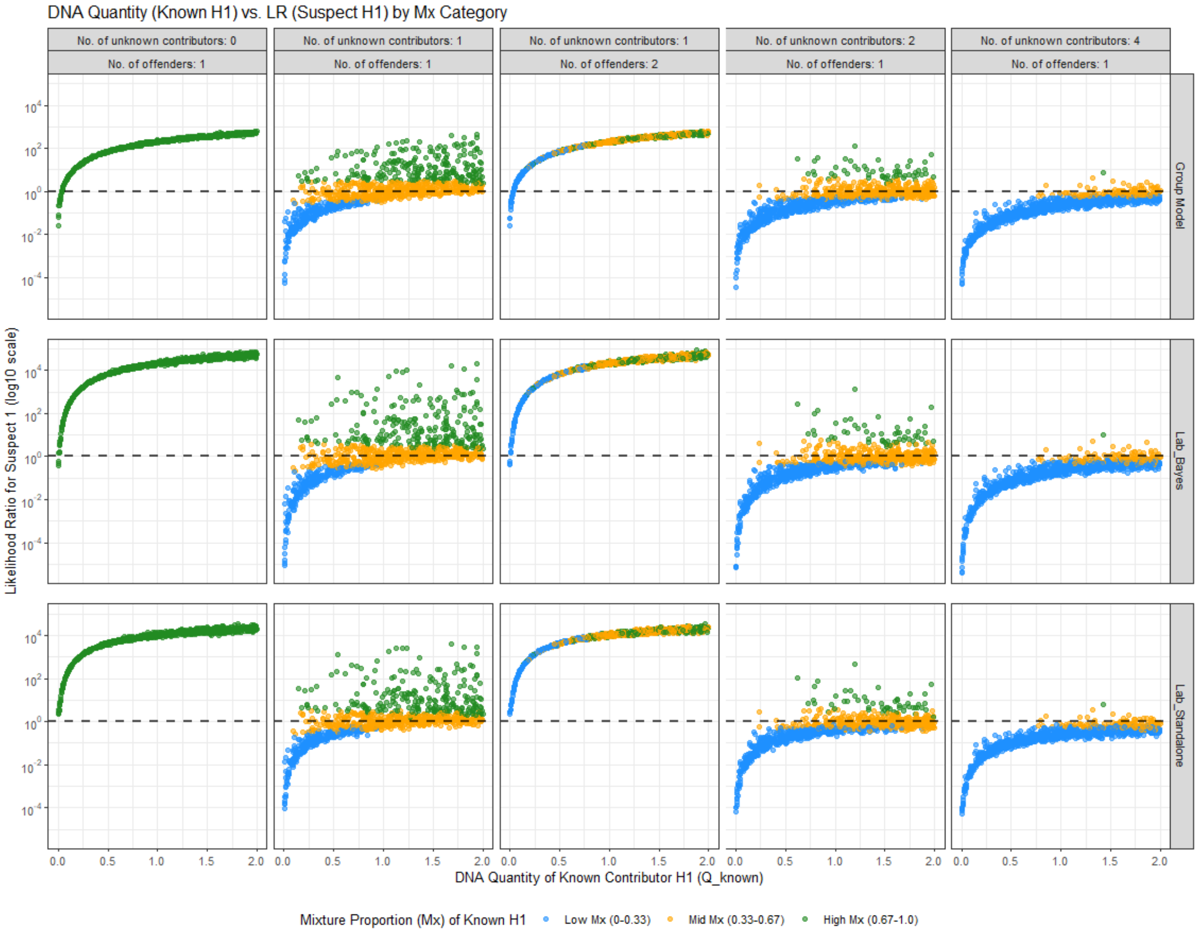


Figure S1: Single-stain simulations (1,000 runs per panel) for `lab_1_ESS`. Columns vary the number of unknown contributors ($N_U$) and assumed relevant actors ($N_S$); rows correspond to the *Group*, *Lab–Bayes*, and *Lab–Vague* models. Each point shows the median LR for S1 versus the realised quantity $q_{S1}$, with colours indicating the realised mixture proportion of S1 ($M_x$). The dashed horizontal line denotes LR = 1.

Several general patterns are evident. When $N_U = 0$ and $N_S = 1$, the LR increases monotonically with $q_{S1}$, with diminishing returns at higher quantities. Introducing unknown contributors ($N_U \geq 1$) spreads the LR over a band, reflecting competition between contributors: high quantities for unknowns reduce the LR for S1, while low quantities increase it. As $N_U$ increases, probability mass is distributed across more potential perpetrators, lowering the typical LR for S1.

The assumed number of relevant actors, assigned here as offenders, plays a critical role. When $N_U = 1$ but $N_S = 2$, the LR collapses back to a narrow curve because the hypothesis that both S1 and the unknown are relevant actors becomes admissible, removing the penalty associated with treating the unknown solely as an alternative to S1.

Across all settings, the *Group* model yields the most compressed and conservative LRs. The *Lab–Bayes* model shows stable and moderated behaviour due to hierarchical shrinkage, while the *Lab–Vague* model exhibits the widest dispersion and occasional tail inflation, particularly at high quantities, due to the absence of cross-laboratory regularisation.

This figure is intended as an illustrative complement to the main results, making explicit that the LR is not a function of DNA quantity alone but depends on the full case context, including contributor structure, relevant actor assumptions, and the underlying model hierarchy.
